# A DNA-based molecular clamp for probing protein interactions and structure under force

**DOI:** 10.1101/2024.06.06.597759

**Authors:** Minhwan Chung, Kun Zhou, John Powell, Chenxiang Lin, Martin A. Schwartz

## Abstract

Cellular mechanotransduction, a process central to cell biology, embryogenesis, adult physiology and multiple diseases, is thought to be mediated by force-driven changes in protein conformation that control protein function. However, methods to study proteins under defined mechanical loads on a biochemical scale are lacking. We report the development of a DNA based device in which the transition between single-stranded and double-stranded DNA applies tension to an attached protein. Using a fragment of the talin rod domain as a test case, negative-stain electron microscopy reveals programmable extension while pull down assays show tension-induced binding to two ligands, ARPC5L and vinculin, known to bind to cryptic sites inside the talin structure. These results demonstrate the utility of the DNA clamp for biochemical studies and potential structural analysis.

## Introduction

Mechanotransduction is now widely recognized as a major determinant of embryonic development, adult physiology and contributor to multiple diseases including cancer, fibrosis, atherosclerosis, osteoporosis and hypertension among others [1, 2]. Cells sense applied forces and mechanical properties of their environment to regulate gene expression and cell behavior across a vast range of biological phenomena.

The major paradigm for cell responses to mechanical variables is that forces across proteins alter their folding landscape to induce conformational changes that modulate enzyme activity or protein-protein interactions [3-5]. In this way, physical forces are converted to biochemical signals that govern cell functions. However, available methods for studying these processes at the molecular level are limited. Single molecular techniques such as magnetic beads or laser traps have greatly advanced the field by measuring effects of force on protein folding and unfolding but are not suitable for analysis of structure or biochemistry [6-8]. Computational methods such as steered molecular dynamics simulations can calculate effects of forces on proteins from first principles and have grown increasingly powerful with improvements in computer speeds [9] . But methods to apply defined forces to proteins in a manner compatible with biochemistry or structural analysis to identify new interactions and confirm computational predictions have not been reported. We therefore set out to develop such a method.

Integrin-mediated adhesions are an important and well-studied locus of mechanotransduction. These mechanical load-bearing structures connect the extracellular matrix to contractile actin filaments inside cells through a set of linker proteins, among which talin1 is central [10, 11]. The talin1 head domain binds directly to β integrin cytoplasmic domains while the rod domain binds both directly to F-actin and indirectly via vinculin. The rod domain is comprised of a series of 4- and 5-α helical bundles that function as springs, unfolding under applied tension and re-folding when tension decreases. These helix bundles bind distinct proteins in their high tension, unfolded state compared to their low tension, folded state, thus serving as force-dependent switches. For example, RIAM1 binds only to folded domains, whereas vinculin binds to sites that become available only when helix bundles unfold under force. Tension-induced vinculin binding then provides additional links to F-actin to strengthen adhesions and resist applied forces. Talin rod domain helix bundles thus provide a well characterized system for validating methods for force application.

We recently identified a new tension-dependent interaction between talin1 and ARPC5L [12], a subunit of the Arp2/3 complex that mediates actin polymerization. This interaction is mediated by the second rod domain helix bundle, R2, which in the complete talin structure exists as a side-to-side dimer with R1. Our results suggest that tension across this dimer opens the R1-R2 interface to reveal the ARPC5L binding site on R2. This event likely occurs before unfolding of the R2 helix bundle that allows vinculin binding. We therefore chose the R1-R2 dimer as our test case for studying protein interactions under force.

## Results

To apply tension to the protein, we developed a method that uses the mechanical properties of DNA, which we have termed the “DNA clamp” (**Fig. 1a**), inspired by studies that utilized DNA to mechanically regulate enzyme activity [13] and employed T4 polymerase to manipulate the shape of DNA origami structures [14]. For this method, a protein with flanking SNAP and Halo tags is reacted with benzylguanine- and chloroalkane oligonucleotide “handles”; these handles hybridize to complementary sequences at the ends of a longer single-stranded DNA (“bridge ssDNA”) that then interconnects the anchored oligonucleotides to form a loop (**Fig 1b**). Conversion of the flexible ssDNA to stiff double-stranded DNA (dsDNA) using DNA polymerase and ligase applies force to the protein.

**Figure 1.**
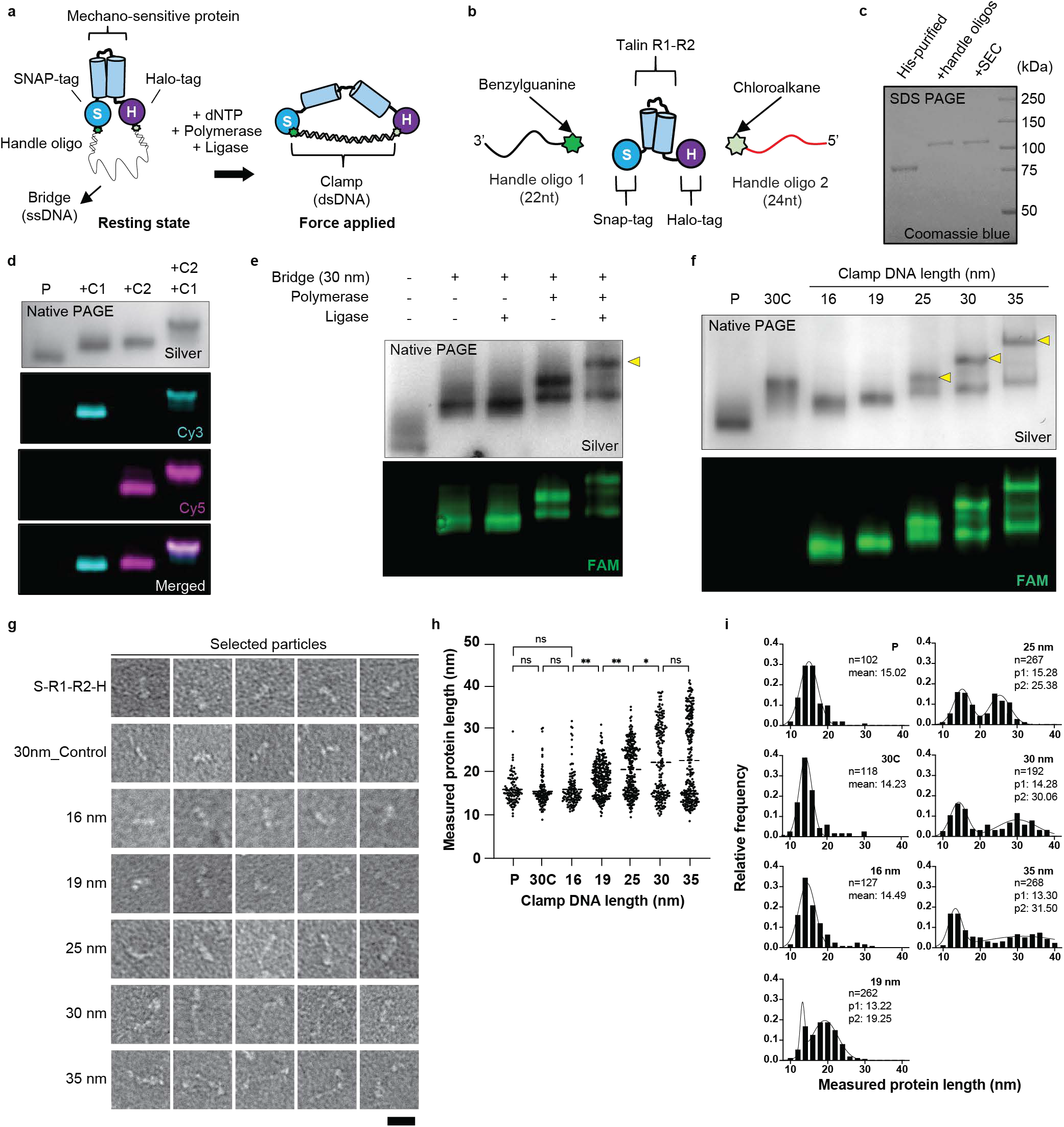
DNA clamp-induced protein conformational change. **a**, Diagram of the DNA clamp design and operation. **b**, Schematics depicting the conjugation reactants: benzylguanine- and chloroalkane-labeled DNA handles and the target protein, talin R1-R2 with terminal SNAP- and Halo-Tags (S-R1-R2-H). **c**, SDS-PAGE showing purified S-R1-R2-H before and after conjugation with DNA handles, with and without purification by size exclusion chromatography (SEC). **d**, Native-PAGE visualizing the handles with fluorescently tagged complementary strands. **e**, Native-PAGE demonstrating band shift after each step of DNA clamp assembly and extension. The 30-nm bridge is fluorescently labeled with Fluorescein (FAM). The fully extended protein band is indicated with a yellow arrowhead. **f**, Band shift observed with DNA clamps of different lengths on native-PAGE, visualized with sliver stain (top) and fluorescence (bottom). Extended protein bands are indicated with yellow arrowheads. **g**, TEM images of the protein with varying DNA clamp lengths. Selected particles represent extended protein for each clamp. Scale bar: 20 nm. **h**, Quantification of the end-to-end distance of the imaged particles by TEM. The means of each group were compared using a one-way ANOVA followed by Tukey’s multiple comparison test (*, P <0.05; **, P<0.01; ns, not significant). **i**, Frequency distribution of the protein end-to-end length for each DNA clamp length, fitted by Gaussian distribution. The histograms of bridge lengths over 19 nm were fitted by the sum of two Gaussian distributions. The total number of measurements and the mean of each Gaussian distribution are indicated in each histogram. Each histogram bar represents a bin range of 2 μm; clamp length is indicated in each histogram. P: protein without the bridge. 30C: protein with a 30 nm bridge lacking the complementary sequence to bind the DNA handle on SNAP-tag side.

We made a construct in which the talin rod domain R1-R2 fragment was engineered with flanking SNAP and Halo tags on either end (**Fig. 1b**). The resultant protein (S-R1-R2-H, or R1-R2 for short) was expressed with a 6×His-tag, purified with Ni-NTA beads, and reacted with a 22-nt benzylguanine-labeled handle and a 24-nt chlorohexane labeled handle. The protein was separated from unreacted oligonucleotides by another round of Ni-NTA purification followed by size exclusion chromatography (SEC, **Fig. 1c**). Conjugation efficiency was assessed by hybridizing Cy5- and Cy3-labeled anti-handles (**Fig. 1d**). The resultant protein contained both fluorophores, confirming successful attachment of the two handles to an R1-R2 protein.

The next step was to connect the two handles at the N- and C-termini of R1-R2 with the bridge ssDNA (**Fig. 1a**). Initially, we used an 88-mer oligonucleotide (dubbed 30 nm bridge for its theoretical length after conversion to dsDNA). The bridge strands were labeled with 6-carboxyfluorescein (FAM) for visualization. Analysis by low-percentage native PAGE revealed a distinct upward shift in the protein band upon bridging (**Fig. 1e**), indicating an increase in Stokes radius. We next added DNA polymerase plus nucleotides to convert the single-stranded bridge to double-stranded DNA. This resulted in a further upward shift. Lastly, we found that including DNA ligase in the system induced an additional small shift, suggesting that sealing the nicks of the newly synthesized DNA provided some further stability or stiffness (**Fig 1e**).

We next used this assay to test bridge ssDNAs of variable length. Longer bridge strands gave rise to greater gel shifts, as expected (**Fig. 1f**). However, with longer DNA strands, a population of unshifted material appeared, suggesting that polymerization efficiency decreased as tension increased. As a control, we also introduced an R1-R2 protein containing only 1 DNA handle at one terminus, which hybridized with the bridge strand but did not complete the loop (30C). After polymerization and ligation, its gel shift was notably less than the dual-handle attached R1-R2 with the same clamp length (30 nm), suggesting that protein conformation as well as DNA clamp length contributes to the gel shift. Together these data validate the assembly of the protein-conjugated DNA clamp.

We next visualized protein conformational changes using negative-stain transmission electron microscopy (TEM). TEM images confirmed extension of the protein that increased with longer clamp length (**Fig. 1g and Fig. S1**). The polymerase and ligase added to the constructs produced negligible background in TEM images (**Fig. S1**). Average protein end-to-end distance increased as expected (**Fig. 1h**). However, as clamp length increased, we also observed broader length distributions and an increase in the population of unextended proteins, more noticeably at 25 nm and above (**Fig. 1i**). These results demonstrate application of force to the protein but also show some structural heterogeneity that increases with degree of extension. This effect may be due to inhibition of T4 polymerase resulting in incomplete polymerization/conversion to dsDNA or kinking of the DNA strand as it nears its persistent length [15], both coupled with the greater folding force generated by the progressively extended protein.

To enrich the population of correctly extended protein, we eluted the upper bands from native polyacrylamide gels of 25, 30 and 35 nm clamp conditions and examined the extracted species by a second band shift assay and by TEM. The native PAGE band shift assay showed enrichment of the slowest migrating bands after the purification (**Fig. S2a**) while quantification of the protein length by TEM demonstrated a higher fraction of extended R1-R2 at the designed clamp length (**Fig. S2b and c**). Purification of the upper band thus improved the results but also suggest some degree of interconversion between fully extended and less extended protein conformations. This implies that the folding energy of R1-R2 approaches the bending energy of DNA clamp when the clamps are longer than 25 nm.

Lastly, we investigated the utility of this approach for biochemical applications. As explained above, the talin R1-R2 region can bind ARPC5L after separation of the folded R1 and R2 domains [12]. It also binds vinculin after unfolding and exposure of the hydrophobic core of either domain [16]. The isolated vinculin head domain D1 is well suited for assaying exposure of buried vinculin binding sites (VBSs) [17]. We therefore purified Flag-tagged version of both the D1 domain of vinculin and ARPC5L (VinD1-flag and ARPC5L-flag), which were immobilized on anti-Flag IgG magnetic beads. Assembled Talin R1-R2 DNA clamps of different lengths were incubated with the beads, washed, and analyzed (**Fig 2a**). R1-R2 binding to both VinD1 and ARPC5L increased after extension. Interestingly, binding to ARPC5L was significant at 25 nm extension whereas binding to VinD1 significantly increased only at 30 nm (**Fig. 2b-e**), suggesting that exposing VBSs requires higher stress on talin than separation of the R1 and R2 domains.

**Figure 2.**
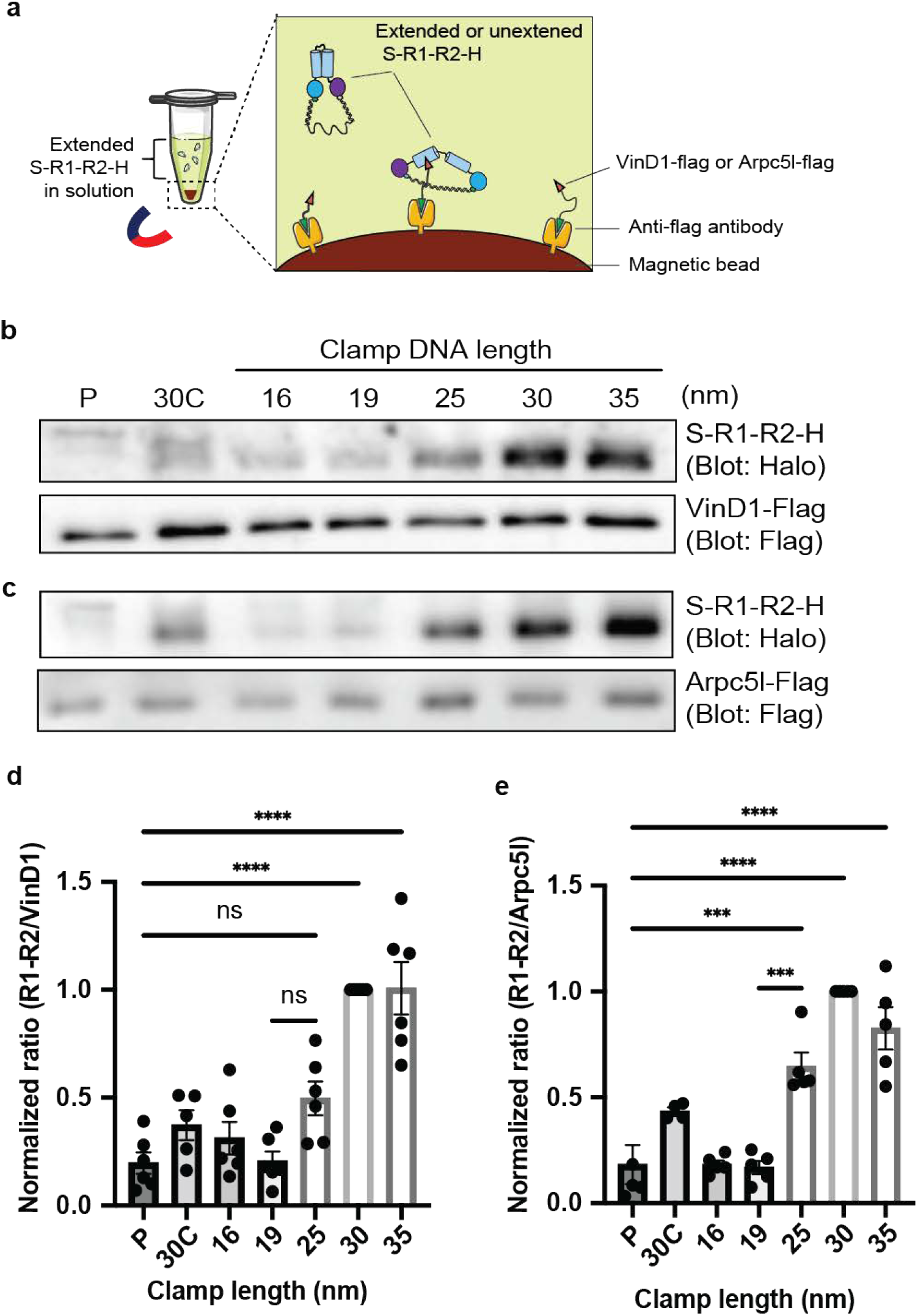
Vinculin and ARPC5L initiate binding to talin at different talin extension lengths. a, Schematic of the pulldown assay. **b and c**, Western blots of S-R1-R2-H with varying clamp lengths pulled down by immobilized VinD1 and ARPC5L, respectively. **d and e**, Quantification of Western blot results. The ratio of S-R1-R2-H to VinD1 or ARPC5L was calculated and normalized to 30 nm condition for each repeat. Group means were compared using a one-way ANOVA (***, P <0.001; ****, P<0.0001; ns, not significant).

## Discussion

We conclude that the DNA clamp can apply forces to proteins resulting in conformational changes and exposure of cryptic binding sites. The device is effective up to extensions in the range of 30 nm, most likely limited by buckling of single DNA duplexes at greater lengths and higher forces. The maximum force that can be generated by our method before buckling of the clamp can be calculated by assuming that DNA behaves as an elastic rod [18]. For DNA clamps of 16, 19, 25, 30 and 35 nm, the maximal forces are 6.9, 4.9, 2.8, 2.0 and 1.4 pN, respectively. Actual force applied to the protein would vary according to the mechanical characteristics and size of the target protein. Note that although the maximum force is smaller than the previously reported force under which these domains fully unfold using magnetic tweezers [19] (R1: ∼22 pN and R2: ∼13 pN), tweezer experiments used high loading rates for much shorter times; application of force for longer times is expected to unfold proteins at much lower forces [20, 21]. Indeed, talin tension reported inside cells by our group and others ranges from 6 to 11 pN [22, 23], although the dynamic range of the FRET sensor used was limited to 11 pN [22]. Our results indicate that under physiological conditions where forces are maintained for longer times, talin domains could be activated at much lower forces than those reported in vitro under high loading rates. The DNA clamp method thus provides a means to identify force-dependent protein interactions. The simplicity in design allows for the control of protein extension length, making it possible to capture multiple force-induced structural states, with or without binding partners. On going work in our labs is aimed at structural studies of proteins under tension and improved methods to apply higher force beyond those achievable with a single DNA duplex.

## Methods

### Protein expression and purification

The Talin 1 rod domain R1-R2 (mouse talin1 482-786) was flanked between SNAP-tag (Addgene #101135) and HaloTag (Promega G8031), followed by sequences encoding a 6xHis-tag. These constructs were then integrated into the pcDNA3.1 vector using Gibson assembly. The Vinculin head D1 domain (1-257, VinD1) and ARPC5L were each expressed in pcDNA3.1 vectors containing C-terminal flag and 6xHis-tags. The plasmids were transfected into 293TX cells using Lipofectamine2000 following the manufacturer’s protocol. After 48 hours, cells were lysed in a phosphate buffer (pH 8.0, containing 300 mM NaCl, 50 mM NaH_2_PO_4_, 15 mM imidazole, 0.2% Tween 20, 1 mM DTT, and EDTA-free protease inhibitor (Pierce). Following brief sonication, the lysate was centrifuged at 30,000g for 30 minutes. The supernatant was then incubated with Ni-NTA beads (Qiagen) for 3 hours with gentle agitation. Beasds were washed with the same buffer and proteins were eluted using 500 mM imidazole. The eluate was subsequently passed through a desalting column (Thermo Scientific 89890, 7K MWCO) pre-equilibrated with PBS (pH 7.4, containing 0.05% Tween 20 and 1 mM DTT) to remove imidazole. The purified proteins were either used immediately for oligonucleotide (oligo) conjugation or snap-frozen for later use.

### Conjugation of oligonucleotides to SNAP and Halo-tag

Benzylguanine (BG, SNAP tag ligand BG-GLA-NHS, NEB, S9151S) and chloroalkane (CL, HaloTag Succinimidyl Ester (O4) Ligand, PROMEGA, P6751) were reconstituted in DMSO to a final concentration of 20 mM. Amino acid-modified DNA oligos (/5AmMC6/ AGGATCGACTAGGTGCGTGGGC and TAGGCAGGTCTGCATACCAGTGAG /3AmMO/ for BG and CL conjugation, respectively) were purchased from IDT and reconstituted in 200 mM HEPES buffer (pH 8.5) to a concentration of 700 µM. The ligand and oligo solutions were mixed in a 1:1 volume ratio and incubated for 4 hours at room temperature. Excess BG and CL labels were removed by ethanol precipitation, and the resulting conjugates were resuspended to a final oligo concentration of 0.5 mM. A four-fold molar excess of the BG and CL-conjugated handles was then added to a 10 µM solution of the purified R1-R2 protein and incubated at 16°C overnight with gentle agitation. Unbound oligos were subsequently removed through Ni-NTA purification followed by size exclusion chromatography using a Superdex 200 column (GE Healthcare). The proteins before and after the reaction were analyzed by SDS-PAGE, stained with Coomassie blue (Thermo Scientific 24615), aliquoted, and stored at -80°C.

### Validation of handle oligos with complementary oligo binding

S-R1-R2-H stock (87.5 kDa, 2 - 4 *μ*M) tagged with handle oligos was reconstituted in 1x T4 ligase buffer (NEB) in PBS with 0.05% NP-40, to achieve a concentration of 0.3 µM or lower. A 1.5x molar excess of Cy3- or Cy5-tagged complementary oligos to the handle oligos was added to the protein and incubated at 37 °C for 1 hour. The resulting mixtures were loaded on a 6% native polyacrylamide gel with SDS-free loading dye (NEB). The gel was made with TAE-Mg buffer (0.4 M Tris, 0.2 M Acetate, 1 mM EDTA, 10 mM MgCl_2_, pH 8.0, 0.05 % NP40) and run in the same buffer for 3 hours at room temperature. The gel was imaged with a laser gel scanner (Typhoon FLA 9000, GE healthcare) to visualize the oligos and then silver stained (Thermo Scientific 24612) to visualize proteins.

### T4 polymerase and ligase treatment for protein extension

A working concentration of T4 nucleotide kinase (PNK, NEB), following the manufacturer’s instructions, and dNTPs (0.2 mM) were added to protein (0.3 µM DNA conjugated S-R1-R2-H protein, 1x T4 ligase buffer in PBS, 0.05% Tween-20). The mixture was aliquoted into PCR tubes and 1.5x molar excess of each of the bridge oligos (5 µM stock in water) were added to each tube and incubated at 37 °C for 2 hours to facilitate annealing. The 5’ 6-carboxyfluorecein (FAM) tagged bridge oligo was purchased from IDT and the list of bridge oligos used in this experiment is provided in the Supplementary Information. In a separate tube, 1 µL of T4 ligase and 0.5 µL of T4 polymerase (both from NEB) were added to 20 µL of PBS. The ligase/polymerase mix was then added to the protein/oligo mix at a portion of 1 µL per 20 µL of the protein/bridge oligo mix. The final mixtures were incubated in a thermocycler following this protocol for the extension reaction: 5 minutes at 25 °C, 3 hours at 16 °C, then ramp-down from 25 °C to 16 °C at a rate of 1 °C per 30 minutes, 1 C stepwise, followed by infinite hold at 12 °C. After the reaction, the resulting gel was examined using the laser gel scanner, and then silver stained. The scale of the reaction was adjusted accordingly.

### TEM imaging and analysis of the extended proteins

The extended S-R1-R2-H proteins were used for TEM (Transmission electron microscope) imaging directly or after purification from a native polyacrylamide gel by excising the desired bands and electroeluting using the MIDI FLEX tube kit (IBI Scientific) in PBS with 10 mM MgCl_2_ and 0.05% Tween 20. The eluted proteins were concentrated using protein concentrator columns (Pierce). Concentration of purified or non-purified proteins were adjusted to 10 – 30 nM. A 5 *μ*L drop of the solution was deposited onto a glow-discharged formvar/carbon-coated copper grid (Electron Microscopy Sciences). The grid was incubated for 2 minutes, and excess fluid was removed by blotting. Grids were rinsed with 5 *μ*L of 2% (w/v) uranyl formate, excess fluid was blotted, and samples stained again with uranyl formate for one minute before blotting and drying. Images were acquired using a JEOL JEM-1400Plus microscope (acceleration voltage: 80 kV) with a bottom-mount 4k×3k CCD camera (Advanced Microscopy Technologies).

### Protein-protein interaction test and western blot

Flag-tagged VinD1 and ARPC5L proteins were immobilized on anti-Flag magnetic beads (MedChemExpress, HY-K0207). A 4 *μ*L suspension of the beads was incubated with 1 - 1.5 *μ*g of extended S-R1-R2-H proteins of varying lengths in PBS (pH 7.4) containing 10 mM MgCl_2_ and 0.1% Tween 20 at 4°C for 1.5 hours with agitation. Beads were washed 3x with the same buffer. Then, the beads were treated with DNase I (NEB) in the manufacturer’s recommended buffer at 37°C with shaking for 30 minutes. Laemmli sample buffer was then added, and samples were incubated at 98°C for 5 minutes. Samples were resolved using SDS-PAGE and transferred to a nitrocellulose membrane using a wet electroblotting system (Bio-Rad). The membrane was blocked using 3% BSA (Millipore Sigma) in TBS with 0.1% Tween 20 (TBS-T) and incubated with the following primary antibodies diluted in TBS-T overnight at 4°C: mouse anti-HaloTag (1:5,000; Promega, G9211), mouse anti-Flag M2 (1:2000; Millipore Sigma, F1804). Membranes were washed with TBS-T for 5 minutes three times on a shaker at room temperature and incubated with HRP-conjugated secondary antibody (1:8000 in TBS-T; Vector Laboratories, PI-2000-1) for one hour at RT, followed by three washes with TBS-T. Bands were visualized using chemiluminescence substrates (Supersignal West Pico Plus or Femto, Thermo Scientific) and imaged on the G:Box system (Syngene).

## Acknowledgments

This work was supported by a National Institutes of Health grant (R01-HL155543) to M.A.S. and C.L., a Department of Defense/Army Research Office MURI grant (W911NF1410403) to M.A.S., a Multidisciplinary Research Initiative Grant W911NF1410403 to M.A.S., a Smith Family Foundation Odyssey Award to C.L., and a fellowship from American Heart Association (20POST35080107) to M.C.

## Figure legends

**Figure S1. TEM images of larger regions at different extension lengths.** In the conditions where the clamp length exceeded 25 nm, correctly extended protein particles are indicated with yellow arrowheads at one end of the particle, while unextended particles are marked with blue arrowheads. Scale bar; 50 nm

**Figure S2. Purification of the upper bands showed reduced unextended population.** a, Native-PAGE showing S-R1-R2-H with various clamp lengths before and after purification. The yellow boxes highlight the excised bands. b, Selected TEM images of the proteins eluted from the excised bands of 25, 30 and 35 nm DNA clamp. Scale bar: 20 nm c, Histogram illustrating the frequency distribution of quantified end-to-end lengths of the proteins, fitted by the sum of two Gaussian distributions. Each histogram bar represents a 2 nm bin range. The mean value and the corresponding clamp length are indicated in each histogram.

