## Supplementary Information for "A DNA-based molecular clamp for probing protein interactions and structure under force"

**List of oligos**

**Handle oligos**

H1: 5’Benzylguanine (BG) tagged, 22 nt

BG-5’AGGATCGACTAGGTGCGTGGGC3’

H2: 3’Chloroalkane (CL) tagged, 24 nt

5’TAGGCAGGTCTGCATACCAGTGAG3’-CL

**Bridge oligos - 6-carboxyfluorecein (FAM) tagged at 5’**
Blue; complementary sequence for H1.

Red; complementary sequence for H2.

**16 nm - 46nt**

5’CTCACTGGTATGCAGACCTGCCTAGCCCACGCACCTAGTCGATCCT3’

**19 nm - 56nt**

5’CTCACTGGTATGCAGACCTGCCTAGAGCCTACCCGCCCACGCACCTAGTCGATCCT3’

**25 nm - 74nt**

5’CTCACTGGTATGCAGACCTGCCTATTCACACTCATTGATATCAGTATAGACTGCCCACGCACCTAGTCGATCCT3’

**30 nm - 88nt**

5’CTCACTGGTATGCAGACCTGCCTATTCACACTCCAGCCTAGAGATATCCCAAGCCTATCATAGACCGCCCACGCACCTAGTCGATCCT3’

**30 nm control - 88nt**

5’TCACCTATGCGTACAGTATCGCATTTCACACTCCAGCCTAGAGATATCCCAAGCCTATCATAGACCGCCCACGCACCTAGTCGATCCT3’

**35nm - 100nt**

5’CTCACTGGTATGCAGACCTGCCTATTCACACTCCACATCAATCCTTGGATATCACTTCAACGAACCCTATCATAGACCGCCCACGCACCTAGTCGATCCT3’
