## Supplementary figures and images for "A DNA-based molecular clamp for probing protein interactions and structure under force"

### Fig. S1

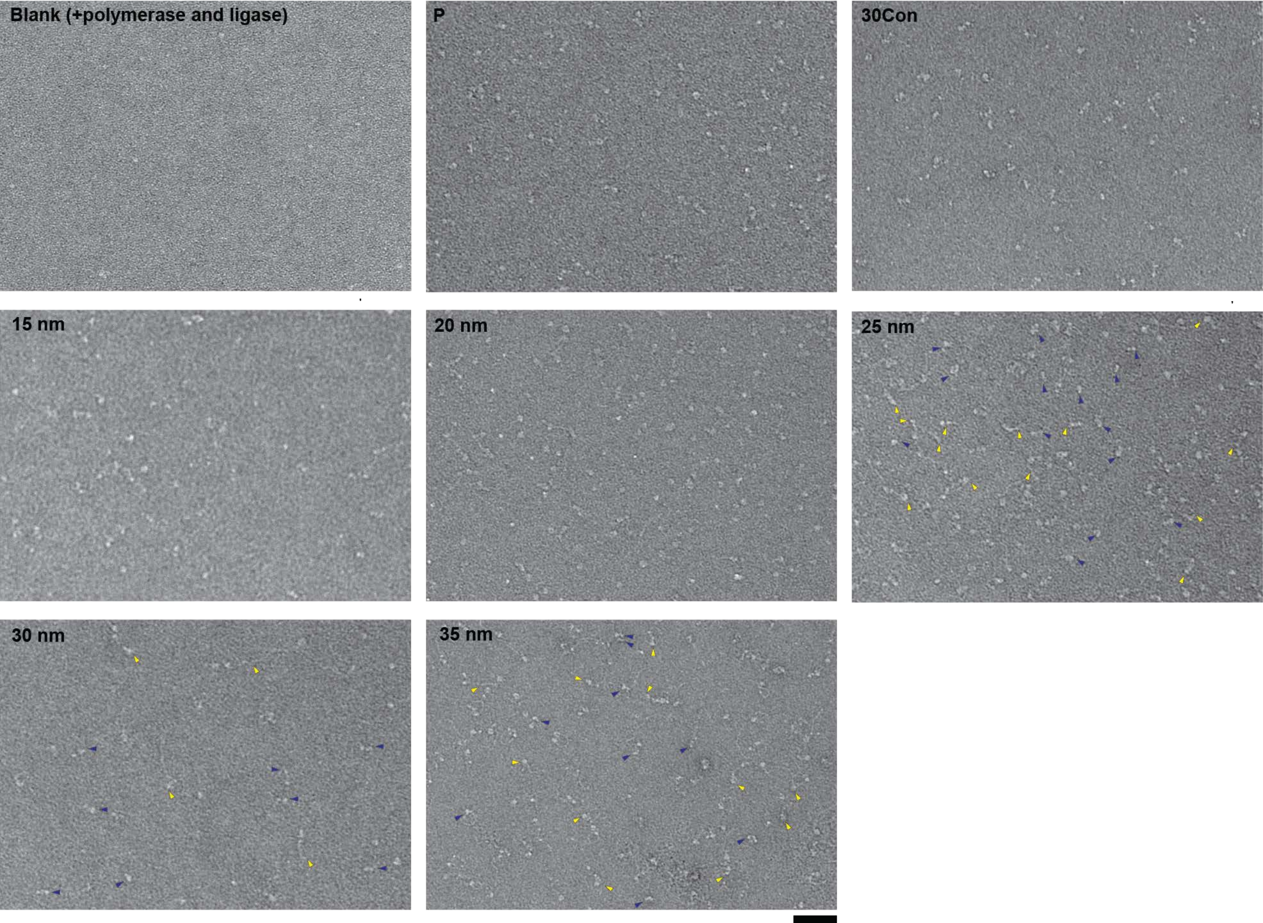

### Fig. S2

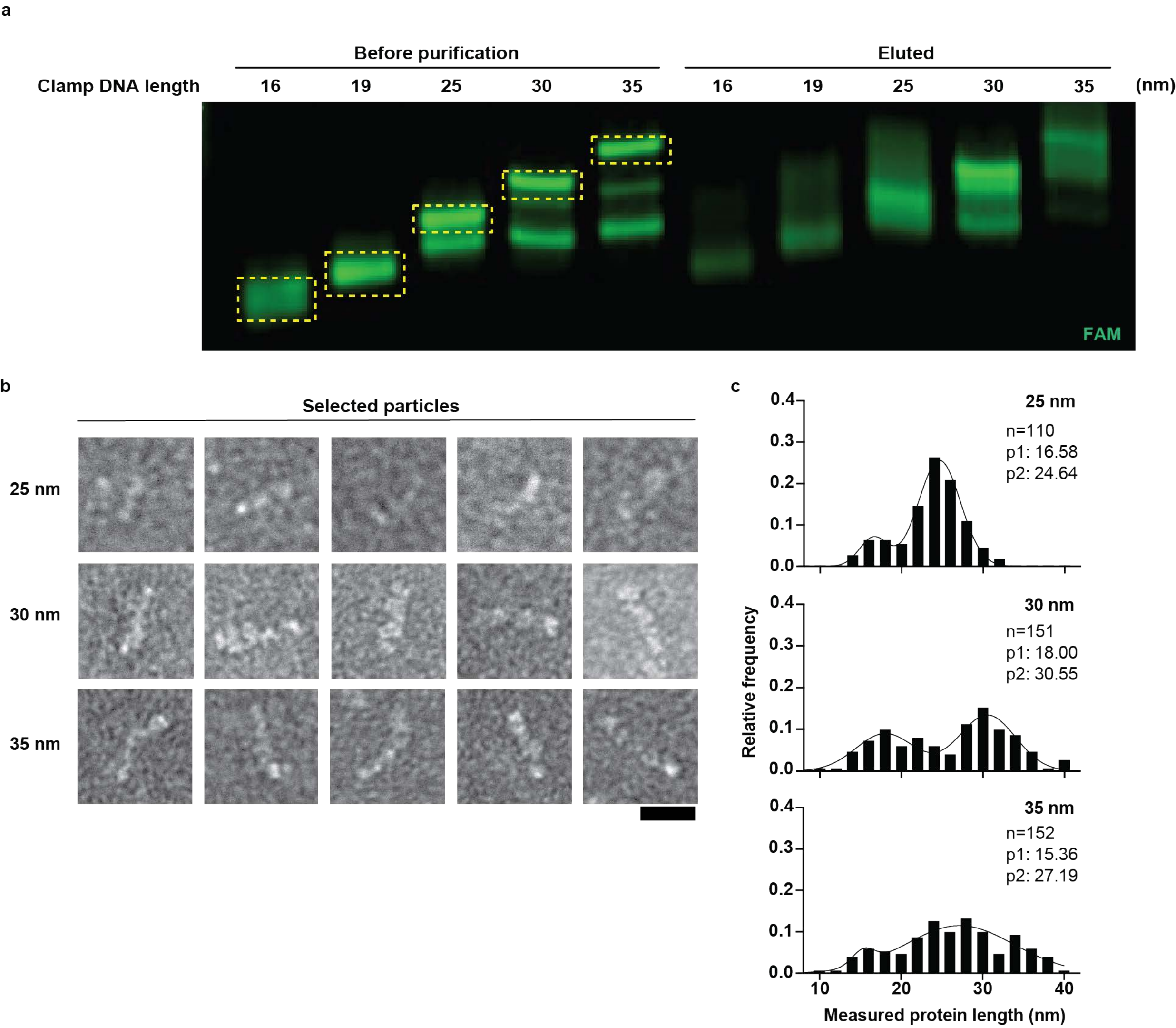
